# Diminished GAD67 Terminals and Ambient GABA Inhibition Associated with Reduced Motor Neuron Recruitment Threshold in the SOD1^G93A^ ALS mouse

**DOI:** 10.1101/2021.12.16.473041

**Authors:** Sharmila Venugopal, Zohal Ghulam-Jhelani, Dwayne Simmons, Scott H Chandler

## Abstract

Pre-symptomatic studies in mouse models of the neurodegenerative motor neuron (MN) disease, Amyotrophic Lateral Sclerosis (ALS) highlight early alterations in intrinsic and synaptic excitability and have supported an excitotoxic theory of MN death. However, a role for synaptic inhibition in disease development is not sufficiently explored among other mechanisms. Since inhibition plays a role in both regulating motor output and in neuroprotection, we examined the age-dependent anatomical changes in inhibitory presynaptic terminals on MN cell bodies using fluorescent immunohistochemistry for GAD67 (GABA) and GlyT2 (glycine) presynaptic proteins comparing ALS-vulnerable trigeminal jaw closer (JC) motor pools with the ALS-resistant extraocular (EO) MNs in the SOD1^G93A^ mouse model for ALS. Our results indicate differential patterns of temporal changes of these terminals in vulnerable versus resilient MNs and relative differences between SOD1^G93A^ and wild-type (WT) MNs. Notably, we found pre-symptomatic up-regulation in inhibitory terminals in the EO MNs while the vulnerable JC MNs mostly showed a decrease in inhibitory terminals. Specifically, there was a statistically significant decrease in the GAD67 somatic abuttal in the SOD1^G93A^ JC MNs compared to WT around P12. Using *in vitro* patch-clamp electrophysiology, we found a parallel decrease in the ambient GABA-dependent tonic inhibition in the SOD1^G93A^ JC MNs. While it is unclear if the two mechanisms are directly related, pharmacological blockade of specific subtype of GABA_A_-α5 receptors suggests that tonic inhibition can control MN recruitment threshold. Furthermore, reduction in tonic GABA current as observed here in the mutant, identifies a putative molecular mechanism explaining our observations of hyperexcitable shifts in JC MN recruitment threshold in the SOD1^G93A^ mouse. Lastly, we showcase non-parametric resampling-based bootstrap statistics for data analyses, and provide the Python code on GitHub for wider reuse.

## Introduction

Amyotrophic lateral sclerosis (ALS) is a neurodegenerative disease that affects motor neurons (MN) of the spinal cord, brainstem, and motor cortex (Cleveland and Rothstein, 2001). The devastating clinical symptoms include weakness, muscle atrophy and fasciculations and rapid progression resulting in fatality within 2–5 years of diagnosis (Hirota et al., 2000). Although MN degeneration is a well-established pathology, not all MNs are equally vulnerable (Kanning et al., 2010). Some MNs such as those controlling ocular and sphincter musculature remain resistant to degeneration (e.g, (Ferrucci et al., 2010; Niessen et al., 2006; Nimchinsky et al., 2000)). Understanding the molecular basis and physiological causes for disease resilience could pave the way for developing effective and early treatments to delay the development and progression of ALS.

One possible basis for disease resistance contributing to neuroprotection is an efficient intracellular Ca^2+^ handling mechanism (von Lewinski and Keller, 2005). Comparative studies further showed a distinct transcriptional profile in resilient oculomotor neurons compared to vulnerable spinal MNs with marked differences in genes with a function in synaptic transmission including several glutamate and GABA receptor subunits (Brockington et al., 2013). This further corresponded with larger post-synaptic GABAergic currents in the resilient ocular MNs compared to vulnerable spinal MNs. ALS-vulnerable brainstem MNs including the trigeminal, hypoglossal, and facial also have differentially distributed GABA and glycine receptors compared with ALS-resistant brainstem MNs (Comley et al., 2015; Lorenzo and Barbe, 2006). Besides these innate differences, in culture and mouse models of ALS, significant alterations have been reported in the two forms of inhibition (glycine and GABA) during the pre-symptomatic stages (Avossa et al., 2006; Chang and Martin, 2009, 2011a, b). However, an equivalent longitudinal examination comparing brainstem vulnerable MNs with resilient ocular MNs has not been reported. Moreover, previously, we reported selective alterations in MN recruitment threshold in vulnerable trigeminal motor pools but not in the resilient oculomotor MNs around postnatal day, P10 (Venugopal et al., 2015). Although changes in membrane excitability have been demonstrated in the vulnerable spinal and hypoglossal MNs at postnatal stages (e.g., (Quinlan et al., 2011; van Zundert et al., 2008)), it remains unclear whether shifts in MN inhibitory mechanisms correspond with, and explain, concomitant shifts in intrinsic MN properties.

To address the above, we conducted comparative analyses of anatomical changes in inhibitory GABA and glycine presynaptic terminals at the motor neuron soma of the well-studied SOD1 mouse model for ALS. Multiple sets of comparisons were made to examine differences between normal (wild-type or WT) and transgenic SOD1 motor neurons further distinguishing ALS-vulnerable trigeminal jaw closer (JC) motor pools and the ALS-resistant extraocular MNs at pre-symptomatic ages (postnatal P12 and adult P50) and symptomatic end stage (P130+) to capture the longitudinal perturbations. Using triple-label fluorescent immunohistochemistry for GAD67 (GABA), glyT2 (glycine) and Choline-acetyl Transferase (ChAT) proteins and established quantitative measures for peri-somatic coverage of GABA and glycine synaptic terminals, we were able to differentiate alterations in these prominent forms of inhibitory terminals based on genotype, age, and disease vulnerability. Secondly, we examined possible changes in ambient GABA-dependent tonic inhibition which can regulate neuronal recruitment threshold (Belelli et al., 2009). Our results provide evidence that tonic inhibition mediated by GABA-α5 receptors controls the recruitment threshold of ALS-vulnerable trigeminal JC MNs. Furthermore, tonic GABAergic current was reduced in the SOD1^G93A^ JC MNs and corroborated with a decreased recruitment threshold in these neurons (Venugopal et al., 2015). In summary, a gradual decline in inhibition appears to a common theme in both vulnerable and resilient MNs during aging which can contribute to excitation/inhibition imbalance and implicated motor and cognitive dysfunctions (Rozycka and Liguz-Lecznar, 2017). Furthermore, the association of decreased tonic GABA inhibition in SOD1G93A JC MNs to the physiological decrement in MN recruitment threshold offer mechanistic insights into emerging muscle dysfunctions.

## Methods

### Transgenic Animals and Genotyping

Transgenic female and male mice overexpressing the human mutant SOD1^G93A^ gene and their non-transgenic wild-type (WT) littermates were used in all of the experiments. All animal protocols were approved by the Institutional Animal Care and Use Committee at UCLA. For genotyping, the hemizygous G93A and wild-type *SOD1* genes were identified using standard PCR techniques (Rosen et al., 1993; Venugopal et al., 2015). Briefly, the mouse tails were cut and incubated in 0.4 ml of lysis buffer (100 mM NaCl, 10 mM Tris, pH 8.0, 25 mM EDTA, 0.5% SDS, 0.1 mg/ml Proteinase K) in a 50°C water bath overnight. After the samples were centrifuged at 6,000 rpm for 10 min, the supernatant was transferred to a tube containing 400 μl of isopropanol. The DNA was collected using flame-sealed capillary pipette and was dissolved in Tris-EDTA buffer, pH 7.6. PCR amplification used the following primers (5’ to 3’): for the wild-type allele, CTAGGCCACAGAATTGAAAGATCT and GTAGGTGGAAATTCTAGCATCATCC, and for the mutant allele, CATCAGCCCTAATCCATCTGA and CGCGACTAACAATCAAAGTGA (Integrated DNA Technologies). The reaction consisted of 30 s at 94°C, 30 s at 57°, and 30 s at 72°C (30 repetitions), and 5 min at 72°C. PCR products were separated on a 2% agarose gel, allowing resolution of a 230-bp product for the wild-type allele and a 194-bp product for the mutant allele (DNeasy 96 Blood and Tissue Kit, Qiagen).

### Animal Groups and Disease Stage Evaluation

For longitudinal evaluation of inhibitory synaptic terminals, we used age-matched WT and SOD1^G93A^ animals at three time points, postnatal age 12±2 days, adult asymptomatic, 50±5 days and 135±10 days corresponding to symptomatic end-stage. For the 3 time points the group sizes were 7, 8 and 12 respectively for WT, and 6, 9 and 11 respectively for SOD1^G93A^ and included both male and female mice. Disease stage was determined by a three-point scoring system to distinguish animals with normal hind limb extension from those at disease onset and end-stage (Herron and Miles, 2012). A score of 2 corresponded to the pre-disease stage with no signs of motor dysfunction. In response to suspension by the base of the tail the hind limbs were symmetrically extended. A score of 1 corresponded to disease onset when the animal started to have hindlimb tremors and/or retraction of one or both hindlimbs when suspended by the tail. A score of 0 corresponded to end-stage of the disease indicated by dragging of at least one hindlimb. For electrophysiology experiments, P10 ± 2 mice of either sex were used.

### Motor neuron electrophysiology

These experiments were conducted on brainstem jaw closer motor neurons located in the dorsolateral region of the motor trigeminal nucleus identified in fresh brain slices *in situ*. Techniques consisted of whole cell current-clamp and voltage-clamp electrophysiology and ion channel pharmacology for GABA receptors. These experiments were conducted in P10 ± 2 day old SOD1^G93A^ mice and their non-transgenic wild-type littermates as controls; Both male and female mice were used.

#### Solutions and Drugs

The brain cutting solution was maintained ice-cold and had the following composition (in mM): 85 NaCl, 2.5 KCl, 1.25 NaH_2_PO_4_, 24 NaHCO_3_, 25 glucose, 75 sucrose, 0.5 CaCl_2_, and 4 MgCl_2_ (Venugopal et al., 2015). The slice incubation solution had the following composition (in mM): 124 NaCl, 3 KCl, 1.25 NaH_2_PO_4_, 26 NaHCO_3_, 10 glucose, 2 CaCl_2_, 2 MgCl_2_, and 4 lactic acid (Del Negro and Chandler, 1997; Hsiao et al., 1998). Current-clamp recordings had a pipette internal solution composition of (in mM) 115 K-gluconate, 9 NaCl, 1 MgCl_2_, 10 HEPES buffer, 0.2 EGTA, 3 K2-ATP, and 1 Na-GTP with a pH between 7.28 and 7.3, and osmolarity between 290 ± 5 mOsm (Venugopal et al., 2015). Voltage-clamp recordings to isolate tonic GABAergic Cl-currents had a pipette internal solution composition of (in mM) 125 CsCl, 5 NaCl, 10 HEPES, 2 MgCl_2_, 0.1 EGTA, 2 Na-ATP, and 0.5 Na-GTP with a pH between 7.28 and 7.3, and osmolarity between 290 ± 5 mOsm (Gupta et al., 2012). AlexaFluor 568 hydrazide (0.2%) was added to the pipette solution for post hoc morphology analysis of recorded neurons. The solutions used in the recording chamber consisted of (in mM concentration), 124 NaCl, 3 KCl, 1.25 NaH_2_PO_4_, 26 NaHCO_3_, 10 glucose, 2 CaCl_2_, and 2 MgCl_2_. In voltage-clamp experiments, tonic GABA current was blocked by GABA_A_-R antagonist 100 μM bicuculline methiodide (Sigma-Aldrich) with control recordings performed in the presence of blockers of excitatory neurotransmission (20 μM DNQX, 20 μM AP5), fast inhibitory neurotransmission (0.5-1 μM Strychnine, 10μM SR95531), and 0.5 μM TTX to block action potentials. GABA-α_5_ receptor inverse agonist L-655-708 (20 μM, Tocris) was bath applied in current-clamp recording to selectively block those receptor subtypes and measure effects on neuron firing properties (Drexler et al., 2013). All drugs and salts were purchased from Sigma-Aldrich unless otherwise stated.

#### Brain slice preparation

Briefly, pups were rapidly decapitated under Isoflurane anesthesia, and the brains were quickly removed and immersed in oxygenated (95% O_2_-5% CO_2_) ice-cold cutting solution. The brainstem was dissected and the rostral end was glued to the platform of the cutting chamber of a vibrating slicer (DSK microslicer, Ted Pella, Redding, CA) and covered with the ice-cold cutting solution. Coronal slices (250 μm thick) were cut and gently placed on a wire-mesh in carboxygenated incubation solution. The slices were incubated at 37°C for 30 min and then maintained at room temperature (22–25°C). To perform electrophysiology recordings from identified JC motor neurons we used retrograde labeling from jaw closer muscles. Briefly, approximately 48 - 72 hours prior to electrophysiology experiments, a retrograde tracer was injected bilaterally (20 μL on each side) into jaw closer muscles of 5-8 day old mouse pups with 1-2% dilution of Alexa Fluor 488 hydrazide (Life Technologies), under isoflurane anesthesia.

#### Whole cell recordings

were performed from trigeminal motor neurons located in the dorsolateral region where the jaw closer motor pools are segregated from the jaw openers (Tanaka et al., 1999). Current and voltage-clamp recordings were performed using the Axopatch 1D patch-clamp amplifier (Axon Instruments, Foster City, CA) and pCLAMP acquisition software (version 9.2, Medical Devices, Sunnyvale, CA). Patch pipettes were fabricated from conventional thin-wall glass (1.5 mm OD, 0.86 mm ID; Warner Instrument, Hamden, CT), pulled on a Brown/Flaming P-97 micropipette puller (Sutter Instruments, Novato, CA) and had bath resistances of 3–5 MΩ. Signals were grounded by a 3 M KCl agar bridge electrode (Ag/AgCl wire) mounted in the recording well. Liquid junction potentials were not corrected. Whole cell capacitance (C_inp_) for each motoneuron recorded in voltage clamp was determined as the time integral of the capacitive current in response to 15-ms, 10 mV voltage command. Uncompensated series resistance (Rs) was calculated from the decay time constant of the transient current and only recordings with <20 MΩ Rs were included for further analysis. The effects of drugs applied to the bath solution were obtained 3–5 min following the onset of drug perfusion. Current-clamp recordings were obtained to study motor neuron firing properties and GABA-α_5_ receptor pharmacology; voltage-clamp recordings were obtained to reveal tonic GABA currents in gap-free mode. Currents measured during voltage-clamp were discarded if they lacked a stable baseline noted as approximately <10% fluctuations in the mean whole cell current. Typically, the giga-ohm seal remained stable for nearly 40-60 minutes following achieving the whole cell mode.

#### Post hoc histology

To visualize neurons using confocal microscopy the dextran, Alexa Fluor 568 was used. The patch pipette was carefully detached from the cell after recording and slices were fixed with 4% paraformaldehyde in phosphate buffer (0.05 M, pH 7.4) for 2–3 days at 4°C. Subsequently, slices are rinsed 2-3 times in PBS for 10-15 mins per rinse and was mounted on a glass slide with coverslip for imaging. All of the procedures were performed at room temperature. A confocal microscope (Zeiss LSM 5) attached to an upright microscope (Zeiss, AxioImager) using Zen Software (Carl Zeiss MicroImaging Inc., Thornwood, NY) was used. The confocal microscope is equipped with single-photon (Argon (488, 514 nm), HeNe (543 nm) and Red Diode (633 nm)) lasers. Z-plane image stacks of each cell were taken at high (63X) and low (10X) magnifications. Optimal Z-plane step sizes were determined on a case-by -case basis with the increment for low magnification always being between 2 and 3 μm. Two-dimensional maximum intensity projection images were then created from the Z-plane image stacks using the Zeiss LSM Image Browser Software and analyzed to confirm cell location and to examine cell morphology.

### Detection and analysis of GAD67 and Gly-T2 presynaptic terminals in juxtaposition to motor neuron soma

Our experimental design and method of quantification of synaptic densities to track anatomical changes in the peri-somatic inhibitory presynaptic terminals on ALS-vulnerable jaw closer (JC) trigeminal and ALS-resistant extraocular (EO) motor neurons, are presented in **Fig. 3** (also see Results). Phenotypically, the transgenic SOD1^G93A^ mice first showed symptoms of unilateral or bilateral hindlimb paresis at P110 ± 4 days which rapidly progressed resulting in weight loss and respiratory failure by P128 ± 15 days, at which point, we regarded as end-stage in this study (Rosen et al., 1993) Whereas, their age-matched wild-type (WT) littermates did not present any such motor deficits and were used as phenotypic and genotypic controls for the various experiments presented.

#### Tissue Preparation

Mice were anesthetized with sodium pentobarbital (2.2 μl /g body weight) and subsequently perfused transcardially with 4% paraformaldehyde (PFA) for 8 minutes and then decapitated. Heads were postfixed overnight with 2% PFA and whole brain region spanning the brainstem/midbrain areas was dissected the following day. The dissected brains were then stored in 30% sucrose overnight. Transverse serial sections (30 μm thick) were cut using a cryostat (Vibratome, Saint Louis, MO) and stored individually in 48-well plates in cryoprotectant buffer (1% polyvinylpyrrolidone, 30% ethylene glycol, and 0.1 mol/L phosphate buffer) at −20°C until used for immunohistochemistry.

#### Immunohistochemistry

For triple-label immunohistochemistry alternate 30μm midbrain/brainstem sections were collected beginning at the level of oculomotor nucleus (Mot III) and extending to the caudal end of the trigeminal nucleus (Mot V). To label motor neurons, the rostral midbrain sections consisting of the oculomotor and trochlear nuclei (Mot III/IV) were stained for ChAT immunofluorescence on the red channel (594 nm), and the caudal pontine sections consisting of Mot V nucleus had retrograde label in the dorsolateral jaw closure motor pools (Venugopal et al., 2015). GABAergic synaptic terminals were detected with antibodies for glutamate decarboxylase 67 (GAD67) (Millipore), and glycinergic synaptic terminals were detected with antibodies for glycine transporter-2 (GlyT2) (Millipore). Sections were stained for GAD67 immunofluorescence on the green channel (488nm) and GlyT2 on the far-red channel (647nm). Briefly, free-floating sections were rinsed in phosphate buffer with saline (PBS) and blocked with 5% normal donkey serum (NDS) diluted with 0.3% Triton X-100 in PBS (TPBS) for 1 hour. Sections were then incubated in primary antibodies diluted with 1% NDS in .3% TPBS for two nights at 37°C. **Table 1** summarizes the primary and secondary antibodies used along with their dilutions. After 3 washes with PBS, sections were incubated for 1 hour at 37°C in a mixture of species-specific secondary antibodies conjugated to Alexa Fluor 488, Alexa Fluor 594, or Alexa Fluor 647 (Invitrogen Corporation, Carlsbad, CA). Sections were washed again in PBS and re-incubated for 1 hour at 37°C in a second mixture of secondary antibodies against the first round of secondaries for further amplification. The tissue was washed with PBS and mounted on frosted glass slides using aqueous media. Triton X (0.3%) was used to increase antibody penetration.

#### Confocal fluorescence imaging

Brainstem sections were first imaged using a 10x water-immersion objective mounted on a Zeiss LSM 5 laser scanning confocal microscope with Zen 2008 software (Carl Zeiss MicroImaging) and subsequently imaged at 63x magnification with an oil-immersion objective using a Zeiss LSM confocal microscope. Three dimensional images were rendered from Z-stacks using Volocity 3D Image Analysis software (PerkerElmer). All images were taken under identical settings and conditions, except for the exposure, which was set for each channel to obtain the best quality image.

#### Motor neuron identification

Choline acetyl transferase (ChAT) immunoreactivity was used as a marker to identify MNs in the oculomotor/trochlear and trigeminal nuclei. For the purposes of this study, it was not possible to unambiguously differentiate oculomotor from trochlear MNs, so we will refer to those eye muscle controlling MNs as a group (Mot III/IV). To demarcate trigeminal motor neurons specifically innervating the jaw closer (JC) muscles a retrograde tracer was injected into the jaw closer muscles; Following 24-48 hours, brains were collected as detailed above and jaw closer (JC) motor neurons were identified in the dorsolateral subdivision in the trigeminal motor nucleus (Venugopal et al., 2015). Neurons labelled by the retrograde tracer showed bright cytosolic puncta in the soma and in most cases, were also co-labeled with ChAT.

#### Detection and quantification of synaptic terminals

Identified oculomotor/trochlear neurons and trigeminal JC motor neurons were analyzed for immunopositive GABAergic and glycinergic synaptic boutons. An average of 3 sections per motor nuclei per animal were included in our analysis. The neuronal numbers analyzed in each experimental group are provided in the respective figures and legends. To control for subjective error during image analyses we ensured that a single researcher conducted this work using the consistent software settings (Volocity). Briefly, the perimeter of a motor neuron’s cell body was traced manually (see **Fig. 3**) and a second contour approximately 2 μm away from the neuron perimeter was drawn to include the synaptic boutons abutting the soma (**Fig. 1B**, dotted white line). Boutons were considered to be in apposition if there was no visible space between the boutons and the neuron’s detected perimeter. Due to the odd shapes of the detected boutons, a Feret’s diameter ≥ 0.18 μm was used as a criterion for inclusion of detected particles (Chang and Martin, 2009). The synaptic abuttal around the soma was calculated as shown in **Fig. 3**. This way, for each MN analyzed, we generated a bivariate dataset consisting of GAD and GlyT-2 synaptic abuttals distinctly.

**Figure 1.**
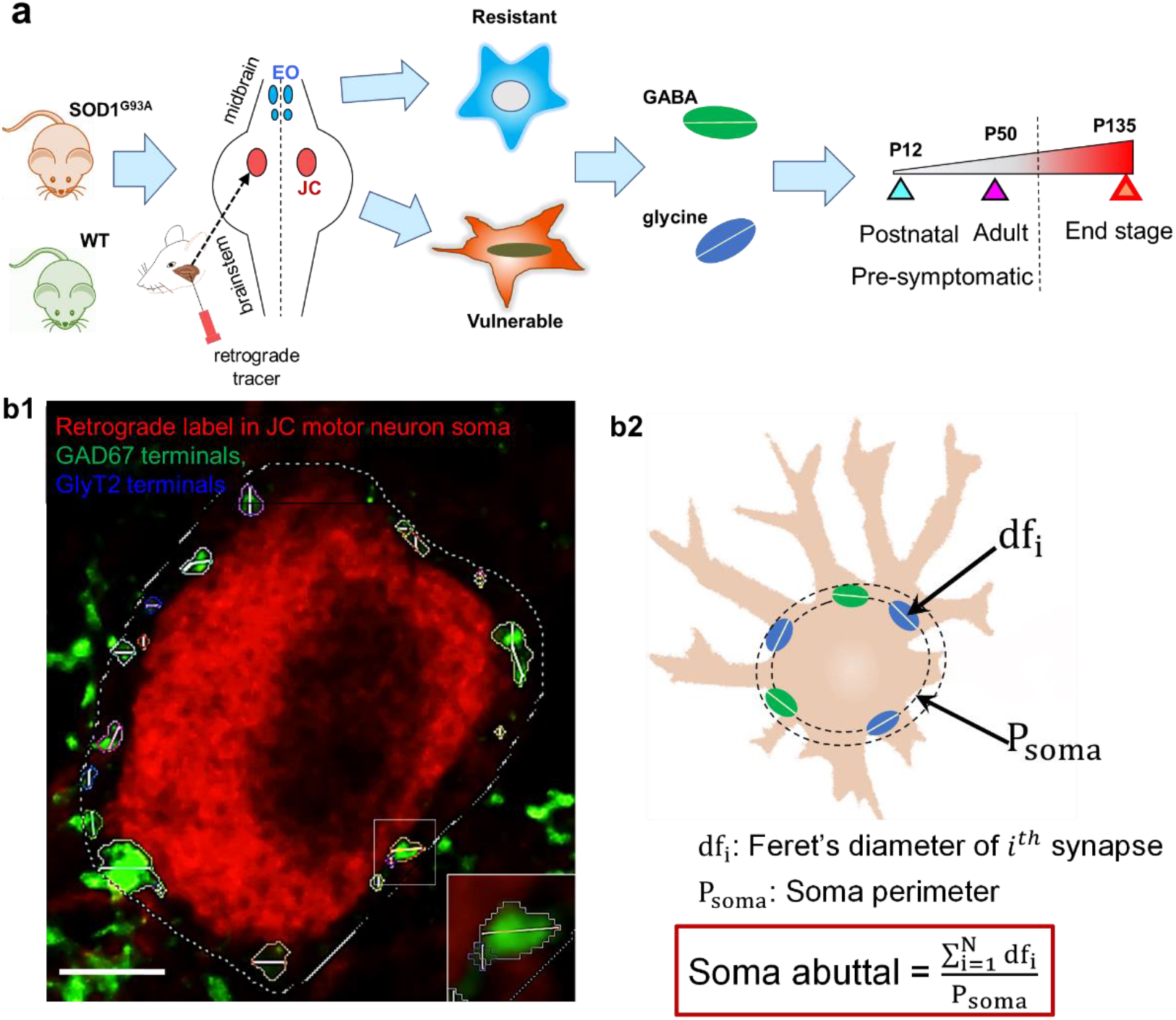
Quantification of GABA and glycine inhibitory synaptic abuttal on motor neuron soma. **a.** Experimental design showing retrograde targeting of ALS-vulnerable jaw closer motor pools and parallel targeting of midbrain. **b1.** Retrograde identification of trigeminal MN (red), GABA (GAD67: green) and Glycine (GlyT2: blue) presynaptic terminals; white dashed contour indicates a user-defined somatic periphery. Synaptic terminals were quantified within this boundary; a single GAD67-positive terminal and a GlyT2-positive terminal are magnified at the bottom right corner highlighting the Ferret’s diameter. **b2.** Schematic showing the quantification method of synaptic abuttal.

**Figure 2.**
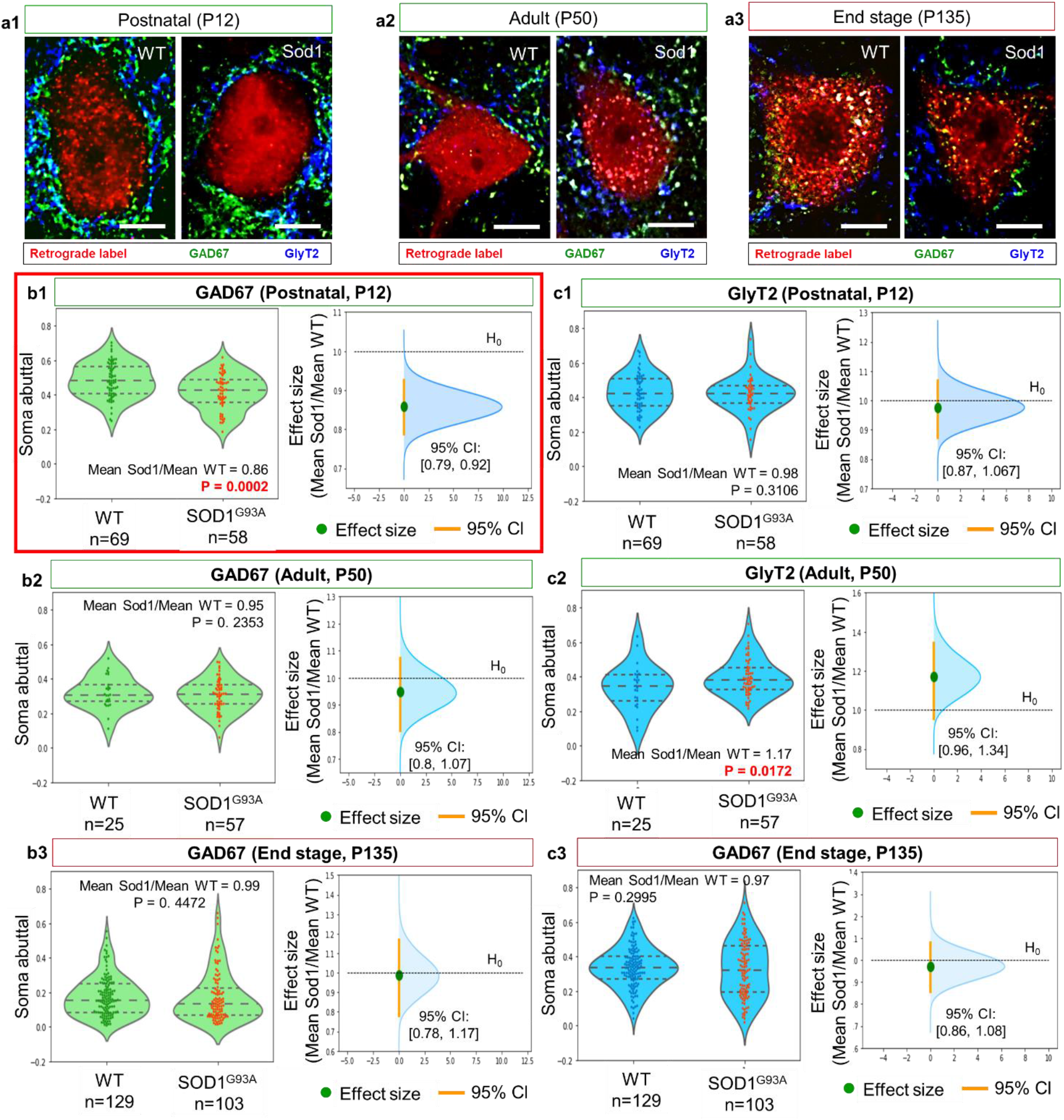
Maladaptive plasticity in GAD67 terminals on ALS-vulnerable motor trigeminal JC MNs in the SOD1^G93A^ mouse. **a1-a3.** Representative images of immunolabeled GAD67 (green) and GlyT2 (blue) terminals around trigeminal motor neurons (red) retrogradely identified by tracer injection into jaw closure muscles. **b1-b3. Left panels:** Violin and beeswarm plots showing WT and SOD1^G93A^ somatic abuttal for GAD67 terminals; **Right panels**: Bootstrap 95% confidence intervals (orange vertical lines) for each pairwise effect size (green circles); blue kernel density histograms show simulated effect sizes for (see resampling Methods). The dashed horizontal line indicates the null ratio of 1. Red box around **b1** indicates statistically significant reduction in peri-somatic GAD67 terminals in the SOD1^G93A^ JC MNs. **c1-c3.** Same descriptors as **b1-b3** for GlyT2 terminals.

**Figure 3.**
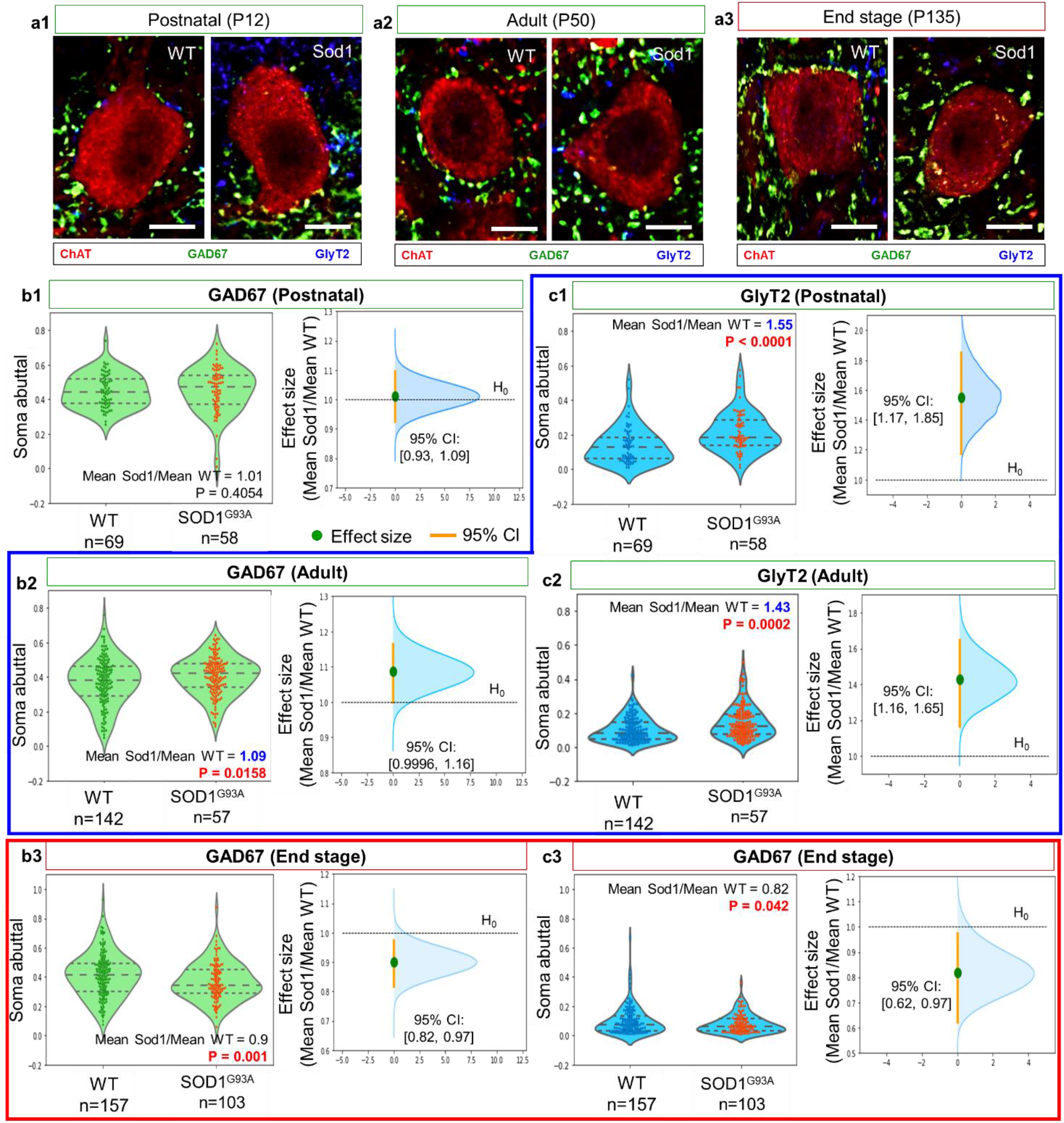
Pre-symptomatic adaptive plasticity in GAD67 and GlyT2 terminals on the ALS-resistant oculomotor neurons in the WT and SOD1^G93A^ mice. a1-a3. Representative images of immunolabeled GAD67 (green) and GlyT2 (blue) terminals around ChAT-stained oculomotor neurons (red). **b1-b3. Left panels:** Violin and beeswarm plots showing WT and SOD1^G93A^ somatic abuttal for GAD67 terminals; **Right panels**: Bootstrap 95% confidence intervals (orange vertical lines) for each pairwise effect size (green circles); blue kernel density histograms show simulated effect sizes from bootstrap resampling (approach in **Methods**). The dashed horizontal line indicates the null ratio of 1. **c1-c3.** Same description as b1-b3 but for GlyT2 terminals. Blue box contours statistically significant *increase* in inhibitory terminals during the pre-symptomatic stages, while the red box contours a statistically significant *decrease* in inhibitory terminals at the symptomatic end stage of the disease.

### Data analysis using non-parametric resampling statistics

We used bootstrap resampling methods to test the Null Hypothesis that there is no difference between groups (e.g., WT vs SOD^1G93A^). Briefly, for two or multiple group comparisons, we conjoined the two or more data sets to be compared into a single set, and resampled with replacement “control” and “treatment” groups from this set. This was repeated 10,000 times to generate the Null distribution, and the 2.5% and 97.5% cutoffs of the 10,000 computed differences were used for an α cutoff of 5% or 0.05 for statistical significance (i.e., p < 0.05). This approach offered freedom of measure and allowed us compare measures beyond just the group mean values (e.g., medians) and effect sizes other than mean differences (e.g., ratios of control and mutant values etc.) (Efron and Tibshirani, 1991). For comparing motor neuron soma synaptic abuttal, we used the ratio of mean SOD^1G93A^ to mean WT values. In place of the classic analysis of variance (ANOVA) which assumes a normal distribution for the underlying populations to compute the F-statistic, we instead computed the F-like statistic as follows:

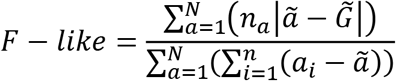

where, *N* is the number of groups being compared, 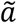 is the group median, 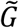 is the grand median of the pooled groups, *a_i_*’s are sample values within each group, and *n* is the group sample size and is different for each group. A p-value was estimated for the computed *F – like* using the procedure described for Null Hypothesis testing above. A pairwise comparison was performed if the *F – like* achieved statistical significance (i.e., p<0.05). To control for false positives, the Benjamini & Hochberg (1995) correction (Hochberg and Benjamini, 1990) was applied for p-values obtained from pairwise comparisons. In addition to p-values, we computed estimation statistics and validate group or pairwise comparisons using 95% confidence intervals (CI) on the effect sizes computed using bootstrap/resampling method (Calmettes et al., 2012). Briefly, to compute the 95% CI, from each sample, repeated pseudo-samples of same size as the original sample were drawn at random, choosing from all the observed values. Each observation remained in the original sample after the value was noted so that values in the original sample could be chosen more than once to make up the pseudo-sample (sampling ‘with replacement’). Thus, each bootstrap sample will usually contained duplicate observations from the original sample. The mean of each pseudo-sample was calculated, and the effect size is computed for the many repeats, similar to the effect size of the original samples. The distribution of these effect size values provided a measure of the confidence limits of the observed effect size (*E_obs_*) of the original samples. For 10,000 such simulated effect size distribution, we used the 249^th^ and the 9749^th^ values as the lower (*E_lower_*) and upper (*E_upper_*) cutoffs respectively, for a 95% CI on the effect sizes. Using the resampling principle (Calmettes et al., 2012), the pivotal 95% CI was obtained as follows:

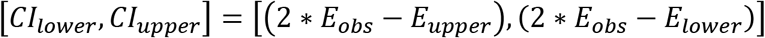

## Results

### Progressive changes in GAD67 and Gly-T2 presynaptic terminals on jaw closer (JC) and extraocular (EO) motor neurons in the SOD1^G93A^ mouse

To compare ongoing homeostatic shifts in inhibitory synaptic terminals in the ALS-vulnerable JC MNs, we undertook an anatomical exploration of both GABA and glycine terminals using presynaptic markers. We further contrasted the abundance of these markers relative to ALS-resistant extraocular motor neurons in the oculomotor and trochlear nuclei (e.g.,(Lorenzo and Barbe, 2006)). Specifically, we measured the peri-somatic coverage of inhibitory presynaptic terminals using immunofluorescence tagged to GAD67 (for GABA) and Gly-T2 (for glycine) synaptic proteins. To track longitudinal changes, three groups of age-matched WT and SOD1^G93A^ mice were included at postnatal (~P12), adult (~P50) and disease end stage (~P135) (see **Fig. 1a**). As shown in the **Fig. 1b1-2**, punctate synaptic densities were detectable around the MN soma, and we quantified somatic abuttal as the ratio of cumulative Feret’s diameter of peri-somatic synaptic densities and the soma perimeter. Statistical comparisons of somatic abuttal between WT and SOD1^G93A^ motor neurons at each of the three time points were conducted as pairwise two-group comparisons as shown **Fig. 1** for JC MNs and **Fig. 2** for EO MNs.

The shapes of the data distribution were seemingly different between WT and SOD1^G93A^ as noted by the violin plots in **Figs. 1** and **2**. Therefore, we used a non-parametric resampling based statistical comparison which is free from assumptions of the underlying data distributions (Calmettes et al., 2012). Such a bootstrap computational approach further allowed us to compare meaningful effect sizes other than the difference between the means, which may not be an appropriate measure when distributions are non-normal such as in our data. As such, we used the ratio of mean somatic abuttal between transgenic and WT as our effect size. Our results show reduced GAD67 somatic abuttal at P12 which was highly significant in the SOD1^G93A^ JC MNs compared to WT (see **Figs. 2b1-b3**). In contrast, statistically significant differences were not detected at adult pre-symptomatic and symptomatic stages. Moreover, Gly-T2 somatic abuttal between SOD1^G93A^ and WT were also not different at the three time points (see **Figs. 2c1-c3**). These results are summarized by the P-values and the 95% confidence limits provided in the respective figure panels.

Next, our analysis of ALS-resistant EO MNs showed interesting contrasts with the vulnerable JC MNs. For example, the mean soma abuttal ratios showed that both GAD67 and Gly-T2 levels were higher in the SOD1^G93A^ EO MNs compared to WT at the two pre-symptomatic time points of P12 and P50 (see **Fig. 3**). Three out of the four of these pair-wise comparisons were also statistically significant (see **Fig. 3b1, b2, c1 and c2**). However, at the disease end stage (P135), both GAD67 and Gly-T2 levels were lower in the SOD1^G93A^ EO MNs compared to WT (see **Figs. 3b3 and c3**). These results are summarized by the P-values and the 95% confidence limits provided in the respective figure panels.

The progressive age-dependent loss in the peri-somatic inhibitory synaptic terminals further highlight the common trends and some contrasts between vulnerable JC MNs and resistant EO MNs (summarized in **Fig. 4**). One-way factor analysis showed the common trend that there is a statistically significant age-dependent decline in both GAD67 and Gly-T2 abuttal in SOD1^G93A^ MNs. Whereas, with age as the factor, WT MNs primarily showed a statistically significant decline in both markers only from P12 to P50; From P50-P135, such decline was absent in all except the GAD67 case for JC MNs in WT neurons. Next, we note that GAD67 showed comparable expression levels in JC and EO WT MNs at P12, but later declined more dramatically in JC MNs compared to EO MNs (see **Fig. 4a1** red versus orange). In contrast, Gly-T2 was more enriched in the JC MNs compared to EO MNs overall (Lorenzo and Barbe, 2006) (see **Fig. 4a1** blue versus purple). Curiously, there was an age-genotype interaction effect from P50 – P135 for three out of the four comparisons shown. The mean values, standard deviations, F-like statistics for multiple group comparisons and pairwise Benjamini-Hochberg corrected P-values are provided in **Fig. 4b1** for WT and **Fig. 4b2** for SOD1^G93A^.

**Figure 4.**
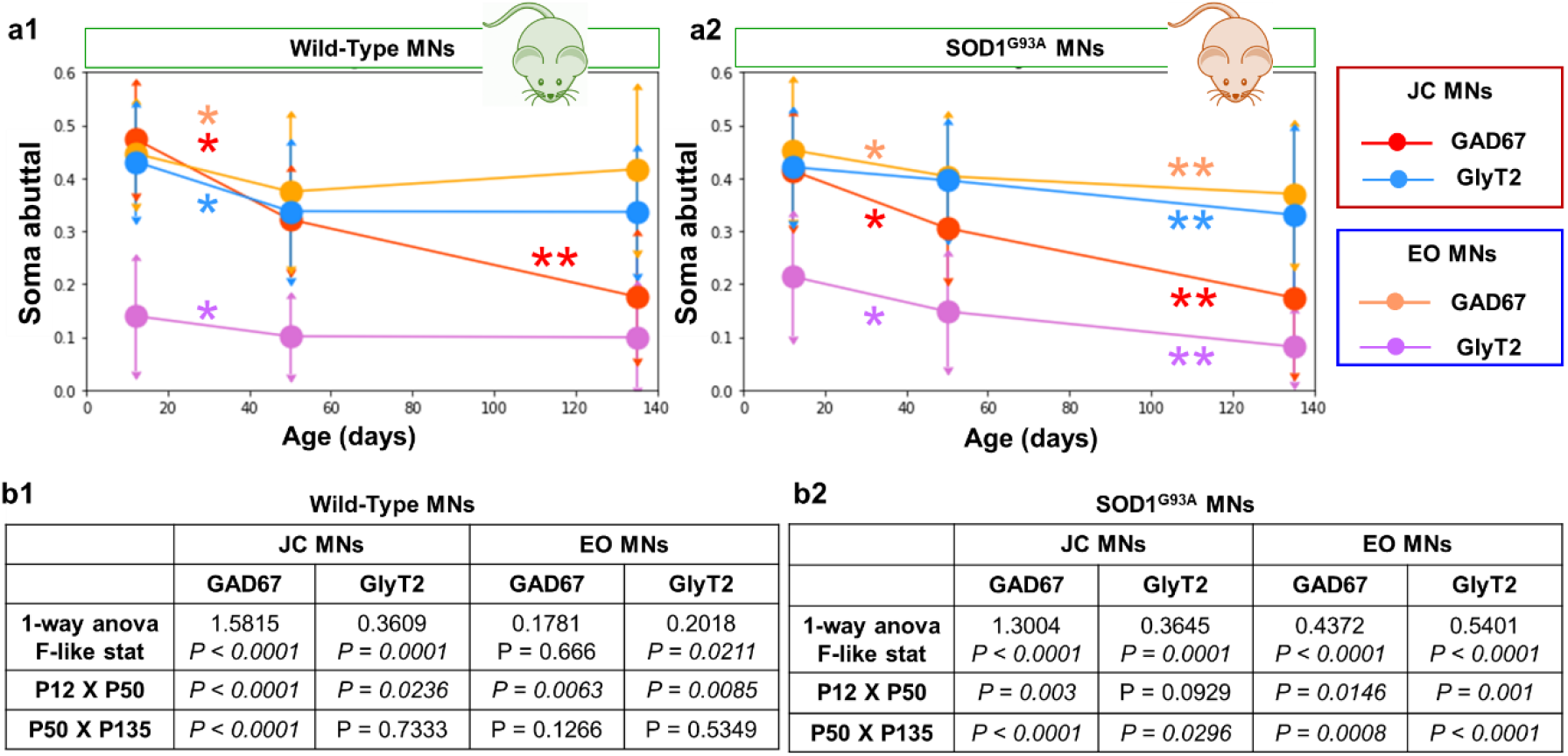
Age-dependent and disease modulated longitudinal alterations in GAD67 and GlyT2 terminals on vulnerable trigeminal JC MNs and resistant EO MNs in the WT and SOD1^G93A^ mice. **a1, a2.** Line graphs show age-dependent changes in synaptic abuttal quantified at pre-symptomatic (P12 and P50) and post-symptomatic end-stage (P135); means (circles) and std. dev. (error bars) are shown for WT **(*a1*)** and SOD1^G93A^ **(*a2*)** mice. The legend on the right shows further grouping based on the type of inhibitory pre-synaptic terminal protein. Single and double asterisks indicate statistical significance at P < 0.05 between age groups. **b1, b2.** Tables show statistics for 1-way ANOVA and pairwise age-group comparisons. All statistical tests were performed using bootstrap resampling approach (see **Methods** and **Results**).

In summary, although early differences existed in GAD67 soma abuttal between WT and SOD1^G93A^ JC MNs only (and not in EO MNs), a decrease in GAD67 levels was striking from the postnatal to adult stage in all of the MNs of both genotypes.

### Reduced tonic GABA inhibition in SOD1^G93A^ JC motor neurons corroborates with hyperexcitable shifts in their recruitment threshold

Given our anatomical finding that GAD67 abuttal is significantly decreased at P12 in the SOD1^G93A^ JC MNs relative to WT, we examined additional inhibitory mechanism which depends on ambient GABA (e.g., (Gupta et al., 2012)). We tested whether the tonic form of physiological GABA receptor current reliant on ambient GABA is reduced in the SOD1^G93A^ JC MNs using *in vitro* whole-cell patch-clamp electrophysiology in retrogradely identifying JC MNs using jaw muscle injections (see **Figs. 5a1 – a3** and **Methods**). In voltage-clamp mode with membrane voltage held at −60 mV as shown in **Figs. 5b1** and **b2**, there was a slow measurable decrease in the holding current from baseline during application of 100 μM Bicuculline Methiodide, a blocker for GABA_A_ receptors. The average shift in holding current density (Δ*I_hold_*) at steady state in the mutant MNs was significantly lower compared to WT (see **Fig. 5b2** for statistical measures). Given that tonic GABA inhibition can regulate MN recruitment current (Torres-Torrelo et al., 2014), the observed reduction of tonic inhibition in the mutant JC MNs could cause a decrease in their recruitment current. Therefore, we conducted a meta-analysis to quantify and compare the recruitment current measured as the soma injected current at the onset of spiking during a triangular ramp current injection in a sample of JC MNs which had showed reduced rheobase current in the mutant (Venugopal et al., 2015), as shown in **Fig. 5c1**. This recruitment current, *I_recruitment_* was significantly smaller for the SOD1^G93A^ JC MNs relative to WT, (see **Fig. 5c2** for statistical measures). To further corroborate the proposition that tonic GABA inhibition regulates MN recruitment, we tested whether blocking specific subtype of GABAΔ-α5 receptors, which mediate tonic inhibition in the spinal neurons (Castro et al., 2011), could indeed reduce the recruitment current in these brainstem JC MNs.

**Figure 5.**
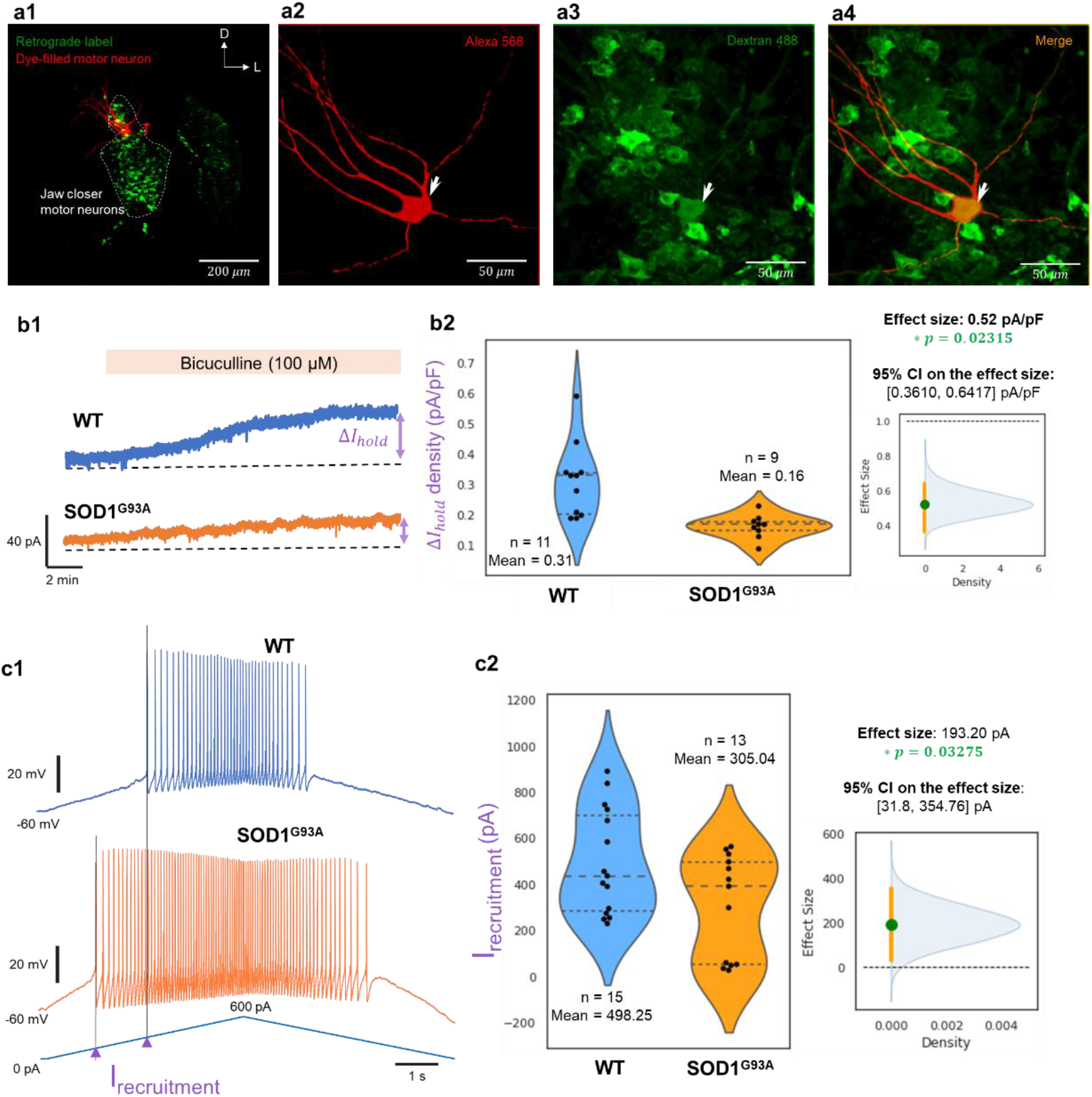
Tonic GABA currents are present and downregulated in the SOD1^G93A^ trigeminal motor neurons at P12±2 days. **a1.** Image shows the dorsolateral trigeminal motor nucleus (dashed contour) with retrogradely labeled motor neurons (green) from jaw closer muscles, while red is a dye-filled patched motor neuron. **a2-a4**. Panels highlight the patched neuron within the labeled nucleus at higher magnification (40x). **b1.** Voltage-clamp recording of whole-cell holding current in gap-free mode in representative WT (blue) and SOD1^G93A^ (orange) JC neurons with membrane potential held at −70 mV. Duration of bath-application of GABA_A_ receptor antagonist bicuculline (100 μM) is marked by the tan rectangle above the current traces. A change in holding current (Δ*I_hold_*) relative to baseline control is highlighted by the purple double arrows. **b2.** Violin plots with overlaid beeswarm plots show holding current density (pA/pF) values at steady-state (at 15-20 mins during Bic application); The n values, effect size as the ratio of mean WT to mean SOD1^G93A^ JC MN current density and two-tailed p-values for two-group comparison is provided. The inset on the right shows the 95% confidence interval (orange vertical lines) for the effect size (green circles); blue kernel density histograms show simulated effect sizes from bootstrap resampling (approach in **Methods**). The dashed horizontal line indicates the null ratio of 1. **c1.** Current-clamp recording of membrane voltage in representative WT (blue) and SOD1^G93A^ (orange) JC neurons during injection of a triangular ramp current (blue lower trace). The spike recruitment current (I_recruitment_) is highlighted by purple arrowheads. **c2.** Violin plots with overlaid beeswarm plots show I_recruitment_ values in putative fast motor units (see **Results** below). The n values, effect size and two-tailed p-values for two-group comparison are provided (also see **Methods** for statistical approach used). The inset on the right shows the 95% confidence interval for the effect size.

We pharmacologically blocked GABA_A_-α5 receptors in normal JC MNs during triangular ramp current injection in current-clamp experiments. As shown in **Fig. 6a1, a2**, bath application of the GABA_A_-α5 receptor inverse agonist, L-655,708 (20 μM) produced a statistically significant decrease in the recruitment current, *I_onset_*. Curiously, as apparent from the interaction plot in **Fig. 6a2**, the decrease in *I_onset_* was greater in MNs with a higher input threshold which form the fast motor units and are preferentially vulnerable in ALS (Kanning et al., 2010). The measured pairwise difference in *I_onset_* or Δ*I_onset_* and the corresponding 95% bootstrap confidence interval for such an observed effect size are shown in the inset in **Fig. 6a2**. Moreover, as shown in the example in panels **a1**, we note that there is indeed deferred termination of firing when GABA_A_-α5 receptors are blocked as noted by a decrease in the injected current at which firing stops (*I_offset_*). Figure panel **6b1** shows the corresponding instantaneous frequency values during the upward ramp (darker colors) and during the downward ramp (lighter colors) in both control (black/grey) and drug (dark/light green) conditions. What is important and interesting to note in these frequency response characteristics is a lack of visible change in the slope of these curves following GABA_A_-α5 receptor blockade. This suggests that tonic inhibition mostly has a shunting effect on MN recruitment and possibly interact with other mechanisms such as L-type Ca^2+^ currents widely present in MNs (ElBasiouny et al., 2010; Hsiao et al., 1998; Venugopal et al., 2011) and which are known to control MN recruitment during muscle force generation (Heckman et al., 2005; Lee et al., 2003). The difference between *I_onset_* and *I_offset_* has been widely used as a physiological correlate of the magnitude of persistent inward currents including the L-type Ca^2+^ and persistent Na^+^ currents in MNs (Hamm et al., 2010; Huh et al., 2017; Jiang et al., 2017; Lee and Heckman, 1998; Powers and Binder, 2001; Turkin et al., 2010). Therefore, we computed the difference between the recruitment, *I_onset_* and de-recruitment, *I_offset_* currents as an estimate of MN persistent inwards currents, and computed Pearson’s correlation with the measured change in the recruitment current due to blockade of tonic GABA inhibition, Δ*I_onset_* as shown in **Fig. 6b2**. There was indeed a very strong correlation which was highly significant as detailed in the figure panel (see statistics in the figure). Together, these results support the co-control of motor neuron recruitment by persistent inward currents and tonic GABAergic inhibition. Given that SOD1 spinal MNs show an increase in L-type Ca^2+^ currents (Quinlan et al., 2011), together with the possibility of reduced tonic inhibition as shown here, we suggest a novel form of imbalance between intrinsic and synaptic regulation of MNs in ALS.

**Figure 6.**
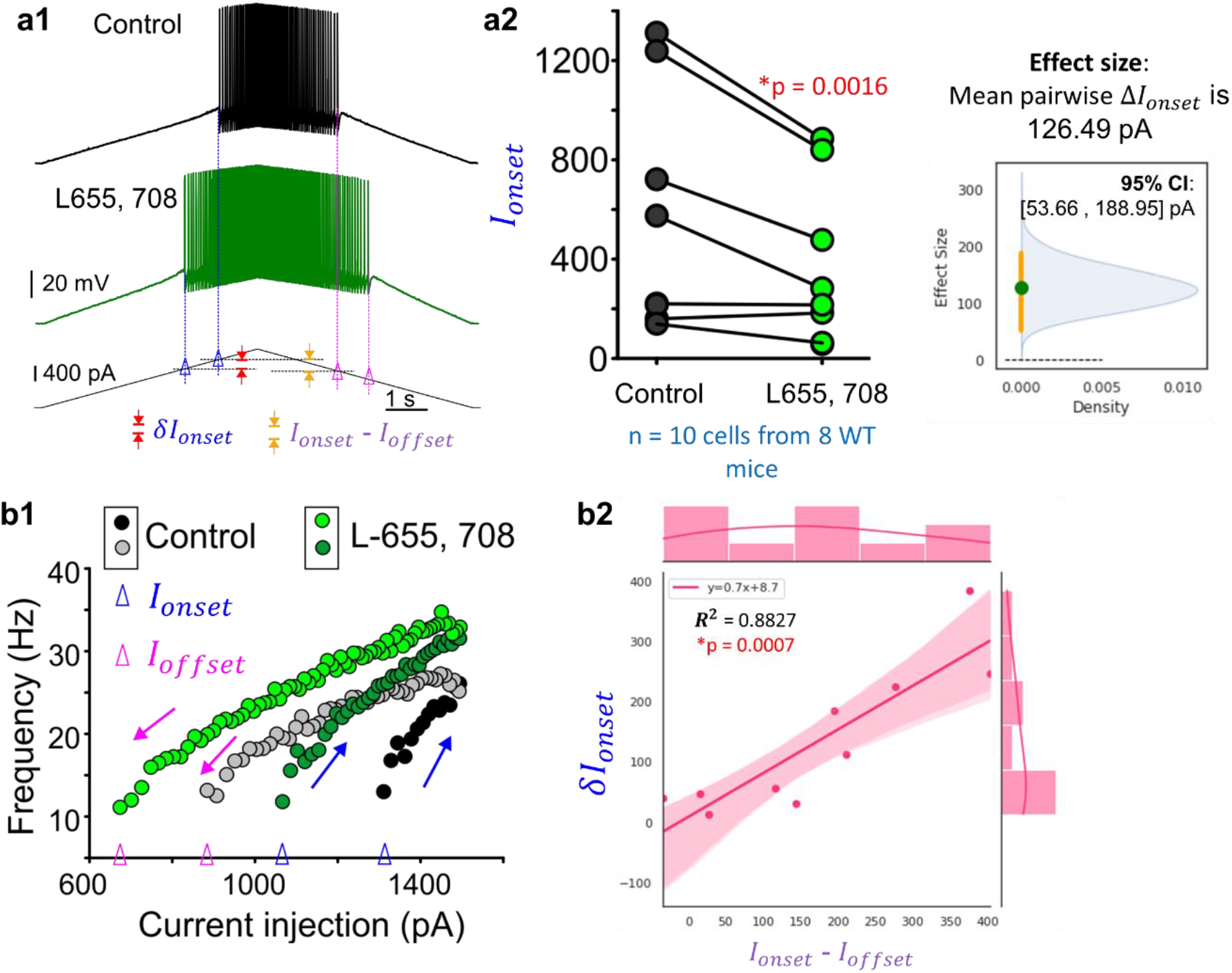
Pharmacological blockade of tonic GABA inhibition in WT motor neurons mimics hyperexcitable shifts observed in the SOD1 trigeminal motor neurons. **a1.** Current clamp recording of membrane voltage during injection of a triangular ramp current (lower trace) in a WT motor neuron. Top black trace shows control condition in nACSF and middle green trace shows response to bath application of 20 μM L-655 708. The spike onset currents under control and drug conditions are marked by blue vertical dashed lines and the open blue triangles on the bottom injected current trace. **a2.** Interaction plot showing significant reductions in spike onset current (Ionset) during L-655, 708 application in WT cells tested. Inset shows the bootstrap 95% CI **b1.** Graph showing the instantaneous spike frequencies as a function of injected current values for the motor neuron presented in **a1**. b2. Linear regression analysis of the change in spike onset current (*δI_onset_*) in relation to an estimate of persistent inward currents as (*I_onset_* - *I_offset_*), where, *I_offset_* is the injected current value at which spike discharge ceases.

## Discussion

In this study we undertook anatomical and physiological examination of inhibitory mechanisms, with the latter focusing on the effects on JC MN recruitment threshold in SOD1^G93A^ mouse and their WT littermates. Our results from immunofluorescent evaluation of GAD67 and GlyT2 labeled presynaptic terminals suggest that both ALS-vulnerable trigeminal JC MNs and ALS-resistance EO MNs of the oculomotor and trochlear motor nuclei show age-dependent decline of inhibition in the SOD1^G93A^ mouse relative to longitudinal changes in the WT neurons. Inhibitory terminals also showed an overall sharp decrease from postnatal to adult stage in both WT and mutant MNs of all the MNs. Interestingly, age-matched comparisons between WT and mutant yielded a statistically significant decrease in GAD67 only at P12 in JC MNs; while such decrease was not apparent in the EO MNs. In contrast to JC MNS, the EO MNs showed an increase in inhibition at both the pre-symptomatic time points of P12 and P50 with statistical significance in 3 out of 4 two-group comparisons (**Fig. 3**). We suggest that such a pattern of increase in inhibition in the resilient EO MNs and not in vulnerable JC MNs may play a role in neuroprotection. For instance, an increase in inhibition could compensate for a parallel disease-induced increase in synaptic excitation (Jiang et al., 2017; Krishnamurthy and Pasinelli, 2021; Krishnamurthy et al., 2021) and in turn maintain homeostasis of excitation/inhibition ratios in the resilient neurons. Whereas, an imbalance between MN excitation and inhibition may underlie a source of vulnerability in the vulnerable JC MNs (e.g., (Schutz, 2005)).

A second curious finding is the existence of a tonic ambient GABA-dependent current in the JC MNs and its decrease at P12 in the mutant corroborating with the anatomical evidence of a decrease in GAD67 terminals. We note that similar to the rat oculomotor neurons, tonic GABA current is likely present in the mouse EO MNs as well, (Torres-Torrelo et al., 2014). However, we did not examine changes in this current in the mutant based on a lack of parallel anatomical evidence. We also did not measure phasic GABA and glycine post-synaptic currents in the JC MNs. While the latter are certainly important for motor neuron outputs (Bae et al., 2002; Bellingham, 2013; Kuo et al., 2003; Li et al., 2004), we focused on the tonic GABA currents which are important for input threshold control in other neurons (e.g., (Belelli et al., 2009; Bonin et al., 2007; Cavelier et al., 2005; Torres-Torrelo et al., 2014)). The key intent was to examine whether our previous observation of reduced recruitment threshold in the SOD1^G93A^ JC MN (Venugopal et al., 2015) could be explained by a reduction in the tonic GABA inhibition. Our results support this possibility since ambient GABA-dependent currents were indeed reduced in the SOD1^G93A^ JC MNs relative to WT (see **Fig. 5**). Moreover, pharmacological blockade of GABAΔ-α5 receptors in normal WT MNs confirmed a role for this current in controlling JC MN recruitment (see **Fig. 6**). Specifically, higher threshold JC MNs (putative fast fatigable motor units) showed a more dramatic decline in input threshold in the presence of L655-708, the inverse agonist of GABA_A_-α5 receptors.

Lastly, we noted that the estimates of the tonic GABA current and that of persistent inward currents (e.g., L-type Ca^2+^ in MNs (Heckman et al., 2005; Hsiao et al., 1998; Hsiao et al., 2005)) from current-clamp in **Fig. 6** show strong correlation. Provided L-type Ca^2+^ currents likely increase as in vulnerable spinal MNs (Quinlan et al., 2011), we suggest that our result showing an association between tonic GABA and L-type Ca^2+^ channels indicates a new form of dys-homeostasis between intrinsic membrane properties and synaptic/extra-synaptic mechanisms (Nieto-Gonzalez et al.). We posit that glial cells which normally tune synaptic excitation/inhibition (Allen, 2014a, b; Ferri et al., 2004; Garnier et al., 2016; Kawamata et al., 2014) and moderate neural excitability (Newman, 2004; Vezzani and Viviani, 2015) and are known to mediate neurodegeneration in ALS (Baufeld et al., 2018; Lasiene and Yamanaka, 2011), may play a role in this excitation/inhibition imbalance in the mutant. These results provide novel directions for identifying early neuroglial markers of synaptic imbalance that could mediate pre-symptomatic dysregulation of neural homeostasis (e.g., (Bataveljic et al., 2012; Kelley et al., 2018)).

## Acknowledgements

This work was supported by NINDS grant NS071348 to SHC, NS095157 to SV and UCLA Academic Senate grant to SHC. We are grateful to Dr. Martina Wiedau, and Dr. Michael Levine Lab members for help with mouse breeding. We thank Dr. Alan Garfinkel for helpful discussions on the resampling statistical methodology. We thank Dr. Xia Yang lab members for providing timely help with molecular analysis.

